# Peak identification and quantification by proteomic mass spectrogram decomposition

**DOI:** 10.1101/2020.08.05.237412

**Authors:** Pasrawin Taechawattananant, Kazuyoshi Yoshii, Yasushi Ishihama

**Affiliations:** Graduate School of Pharmaceutical Sciences, Kyoto University, Kyoto 606-8501, Japan; Graduate School of Informatics, Kyoto University, Kyoto 606-8501, Japan; RIKEN Center for Advanced Intelligence Project (AIP), Tokyo 103-0027, Japan; Laboratory of Clinical and Analytical Chemistry, National Institute of Biomedical Innovation, Health and Nutrition, Ibaraki, Osaka, 567-0085, Japan

## Abstract

Recent advances in liquid chromatography/mass spectrometry (LC/MS) technology have notably improved the sensitivity, resolution, and speed of proteome analysis, resulting in increasing demand for more sophisticated algorithms to interpret complex mass spectrograms. Here, we propose a novel statistical method that we call proteomic mass spectrogram decomposition (ProtMSD) for joint identification and quantification of peptides and proteins. Given the proteomic mass spectrogram and the reference mass spectra of all possible peptide ions associated with proteins as a dictionary, our method directly estimates the temporal intensity curves of those peptide ions, i.e., the chromatograms, under a group sparsity constraint without using the conventional careful pre-processing (e.g., thresholding and peak picking). We show that the accuracy of protein identification was significantly improved by using the protein-peptide hierarchical relationships, the isotopic distribution profiles and predicted retention times of peptide ions and the pre-learned mass spectra of noise. In the analysis of *E. coli* cell lysate, our ProtMSD showed excellent agreement (3277 peptide ions (94.79%) and 493 proteins (98.21%)) with the conventional cascading approach to identification and quantification based on Mascot and Skyline. This is the first attempt to use a matrix decomposition technique as a tool for LC/MS-based joint proteome identification and quantification.

## INTRODUCTION

Liquid chromatography coupled with mass spectrometry (LC/MS) has been widely used to identify and quantify peptides obtained by digestion of proteins in shotgun proteomics. Recent advances in LC/MS technologies have permitted the detection of peaks with high resolution and high accuracy in a two-dimensional mass spectrogram over the time-*m/z*-domain. Despite the high performance of modern LC/MS instruments, however, a large part of the mass spectrogram remains unresolved.^1^ The common data analysis techniques start with pre-processing steps that simplify mass spectrograms at the cost of information loss and error propagation.^2^ A new approach based on mathematical and statistical methods is required to interpret mass spectrograms accurately with higher sensitivity.

The mass spectrogram can be viewed as a nonnegative matrix in which peaks are continuously recorded. The peaks of peptide ions in a mass spectrogram are sparse in general, meaning that a relatively small proportion of interesting signals is present over a large space on the mass spectrogram, and their dynamic range can extend over many orders of magnitude. Multivariate curve resolution-alternating least-squares (MCR-ALS) was recently proposed to analyze high-resolution LC/MS data in a matrix form with a simple sparsity constraint and without using a prior threshold.^3^ The algorithm is well designed for analyzing sparse data and identifying low-intensity *m/z* signals. However, only eight resolved peak profiles from eight chemicals and four peak profiles from bacterial cells were considered. In addition, a pre-processing step of manual selection of chromatographic windows to reduce complexity is required. The analysis results also include negative intensities, which are inconsistent with the nonnegative nature of the mass spectrogram.

In the field of statistical signal processing, nonnegative matrix factorization with a sparsity constraint has been a powerful tool for audio source separation. Specifically, the time-frequency spectrogram of a mixture signal (nonnegative matrix) is approximated by the product of two nonnegative matrices, namely a dictionary and an activation matrix, where the dictionary is (partially) given or estimated. In this paper, data analysis with a fixed dictionary is called *decomposition* instead of *factorization*. For LC/MS data, it was reported that nonnegative matrix decomposition enabled identification of 11 out of 18 mixed chemicals.^4^ To our knowledge, it has not yet been explored whether such a naive application can be used for more complex real-world samples, including proteomic mass spectrograms.

In this study, we propose a new approach that we call proteomic mass spectrogram decomposition (ProtMSD) for joint identification and quantification of peptides and proteins using shotgun proteomic samples. This method incorporates the knowledge of the isotopic *m/z* distributions, the learned mass spectra of noise, and the possible retention times of peptides. In order to enhance the group sparsity in the proteomic data, we focus on the protein-peptide hierarchical relationships. Using a standard sample, we evaluated variants of ProtMSD with and without prior knowledge with a fixed dictionary of all possible peptides. We then applied ProtMSD with a dictionary based on the peptide detectability information for analyzing a large proteome-scale mass spectrogram from an *E. coli* digest. We confirmed that ProtMSD successfully increased the number of identified proteins and improved the accuracy of quantification without the need for complicated pre-processing (e.g., thresholding and peak picking) which carries the risk of information loss.

## EXPERIMENTAL SECTION

### METHOD

We first review nonnegative matrix decomposition and then explain its application to proteomic mass spectrograms.

#### 1. Nonnegative matrix decomposition

The goal of nonnegative matrix decomposition or factorization is to approximate a nonnegative matrix 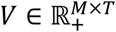 as the product of two nonnegative matrices 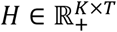 and 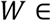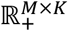 as follows:

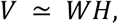

where *K* is the number of basis components. One of the most popular cost functions for evaluating the approximation error between *V* and *WH* is the generalized Kullback-Leibler (KL) divergence given by

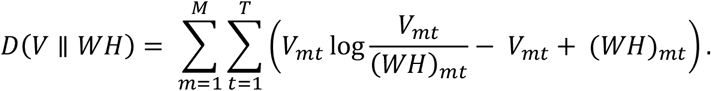

Because optimal *W* and *H* cannot be computed in a closed form, one needs to initialize *W* and *H* and alternately and iteratively update them until convergence is achieved.^5^ When *W* is given and fixed in the decomposition setting, it is still necessary to use an iterative optimization algorithm for estimating *H* because *V* ⁄ *W* is not mathematically defined.

#### 2. Proteomic Mass Spectrogram Decomposition (ProtMSD)

We deal with a two-dimensional mass spectrogram 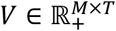 of a shotgun proteomic sample represented in a nonnegative matrix form, in which each element *V*_*mt*_ (1 ≤ *m* ≤ *M*, 1 ≤ *t* ≤ *T*) indicates the nonnegative intensity at *m/z* bin *m* and time frame *t*, where *M* is the number of *m/z* bins and *T* is the number of time frames. To obtain *V*, the intensities of peaks detected in LC/MS measurement are quantified with certain *m/z* and time resolutions. Finally, *V* is normalized such that the mean of *V* is equal to 1.

Our decomposition method, ProtMSD, has three key features that differ from standard matrix decomposition. First, ProtMSD uses a fixed dictionary based on prior knowledge about peptide ions and learned noise components. Assuming that *S* peptides (originating from *G* proteins) and *N* noise components, i.e., *K*_*S*_ + *K*_*N*_ components in total, might be contained in *V*, we aim to estimate an activation matrix 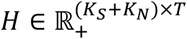 consisting of the temporal intensity curves of those components by referring to a fixed dictionary 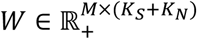 consisting of the mass spectra of those components such that *V* ≃ *WH*. Each column *k* of the signal dictionary *W*_*S*_, (*W*_*S*_)_*k*:_ (1 ≤ *k* ≤ *K*_*S*_), is the theoretical mass spectrum of peptide *k* consisting of isotopic peaks of the possible peptide ions, where “:” represents a set of all indices. Each column *k* of the noise dictionary *W*_*N*_, (*W*_*N*_)_*k*:_ (1 ≤ *k* ≤ *K*_*N*_), is the mass spectrum of a noise component. In advance of ProtMSD, *W*_*N*_ is estimated by using standard matrix decomposition with *K*_*N*_ basis components for mass spectrograms of noise including no peptide ions (e.g., last 10 min of gradient elution). Because mass spectrograms are usually contaminated by considerable fairly constant background noise, ProtMSD based on both the peptide signal and noise dictionaries is expected to accurately and directly quantify peptides from noisy measurement data.^6^

Second, ProtMSD allows each peptide to be activated at time frames in which the peptide is considered highly likely to be detected. More specifically, for each peptide *k* (1 ≤ *k* ≤ *K*_*S*_), the activation (*H*_*S*_)_*kt*_ at frame *t* is allowed to become larger than zero if *t* is around the predicted retention time (given by achrom^8^) or the retention time observed in the reference run. Otherwise, the activation (*H*_*S*_)_*kt*_ is restricted to zero. Thanks to the multiplicative nature of the iterative updating algorithm, the activations at unlikely time frames are kept at zero once they are initialized to zero.

Third, ProtMSD incorporates the group sparsity based on the hierarchical relationships between proteins and peptides. If a protein is (not) included in a target mixture, many peptides obtained by digesting the protein are expected (not) to be detected simultaneously. More specifically, we put a log/*L*1 regularization term^7^ on *H* as Ω(*H*).

Given *V* as an observed mass spectrogram and a fixed dictionary *W*, the goal of ProtMSD is to estimate *H* under the constraint that *D*(*V*||*WH*) + Ω(*H*) is minimized. To achieve this, we derive an iterative optimization algorithm based on the majorization-minimization principle.

#### 3. Identification and Quantification of Peptides and Proteins

We identify and quantify peptides and proteins included in the target mixture from the activation matrix *H* estimated by ProtMSD. First, *H* is multiplied by the mean intensity of the original unnormalized mass spectrogram. Peptide *k* (1 ≤ *k* ≤ *K*) is judged as included if the maximum value of the intensity curve (*H*_*S*_)_*k*:_ of each peptide *k* is 3-times larger than the noise. Protein *g* is reported if at least two peptides corresponding to the protein are identified. The quantity of protein *g* is given as the sum of the intensities of the detected peptides.

Further details of calculation methods, matrix constructions, the protMSD algorithm, and post-processing techniques are provided in Supplementary Methods.

We conducted a proof-of-concept experiment in which ProtMSD is applied to the mass spectrogram of a mixture of peptides obtained by digesting four proteins. We also conducted a proteome-scale experiment using *E. coli* cell lysate data.^8^ Further details are given in the supplementary document. All computations were performed on a Xeon E3-1226 v3 3.30 GHz CPU with a memory of 32 GB. The raw MS data of the four-protein mixture and *E. coli* lysate were registered with identifiers JPST000765/PXD015189 and JPST000663/PXD018074, respectively, at the ProteomeXchange Consortium via jPOST partner repository.^9^ The code of ProtMSD is implemented in Python 2.7 is available at GitHub https://github.com/pasrawin/ProteomicMSD and https://github.com/pasrawin/LibraryProteomicMSD.

## RESULTS AND DISCUSSION

### 1. Application to four-protein digestion standard

#### 1.1 Effectiveness of Prior Knowledge

We validate the effectiveness of the prior knowledge by comparing two variants of ProtMSD with and without isotopic distribution, learned noise dictionary, retention time initialization, and group sparsity constraint, designated as ProtMSD and ProtMSD0, respectively, for convenience. The four-protein digest afforded 1057 peptide ions. We found a proportional relationship between the observed total ion current (TIC) chromatogram (Figure 2A) and the sum of the intensity curves estimated by ProtMSD0 (Figure 2B), with a correlation coefficient of 0.846. On average, 20% of the TIC chromatogram was accounted for. To investigate the effectiveness of precursor ion isotopic distributions, noise models, sparsity constraints, and the other factors, we evaluated the correlation to the observed TIC chromatogram (Figure 2C). With ProtMSD, we obtained a correlation coefficient of 0.982 and could account for 70% of the TIC chromatogram. The remaining signals were due to peptides in undefined charge states, peptides with undefined modifications, contaminants, and non-electronic noise outside the ProtMSD dictionary, because the TIC chromatogram included all signal contributions observed during LC/MS.

**Figure 1.**
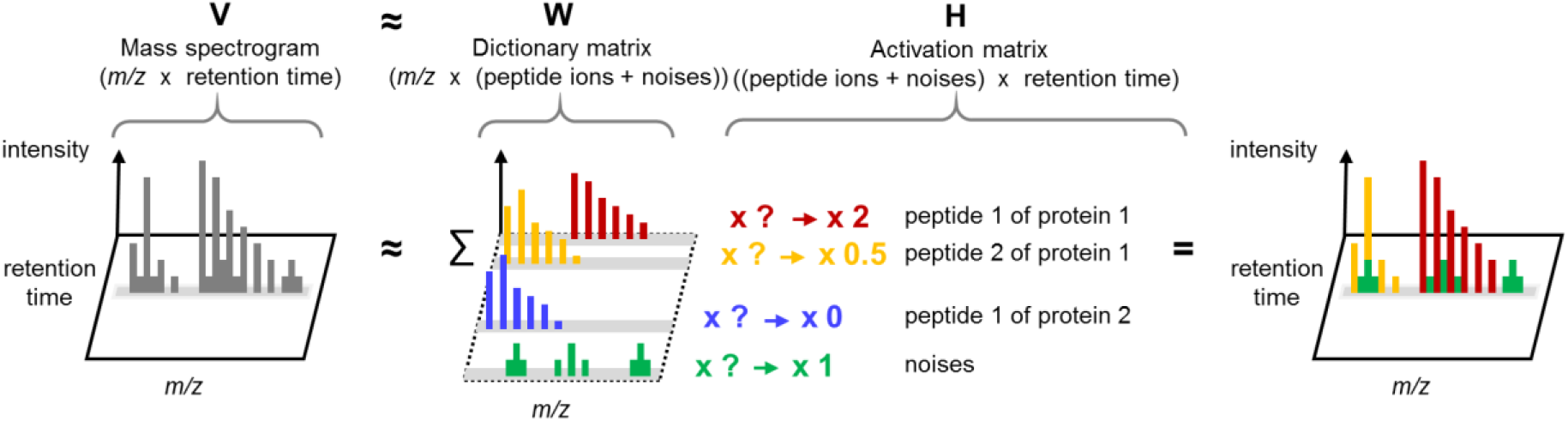
Overview of ProtMSD. Under the linear mixture model, a vector along the *m/z* axis of *V* represents an experimental MS1 spectrum. ProtMSD aims to solve nonnegative values for *H* using the theoretical isotopic distribution of peptide ions in *W* whose product *WH* well approximates *V*. Under the matrix model, it can be written as *V* ≃ *WH*. The group sparsity constraint restricts the approximation to be sparse, and the peptide signal and noise dictionaries help in separating peptides from noises for quantification.

**Figure 2.**
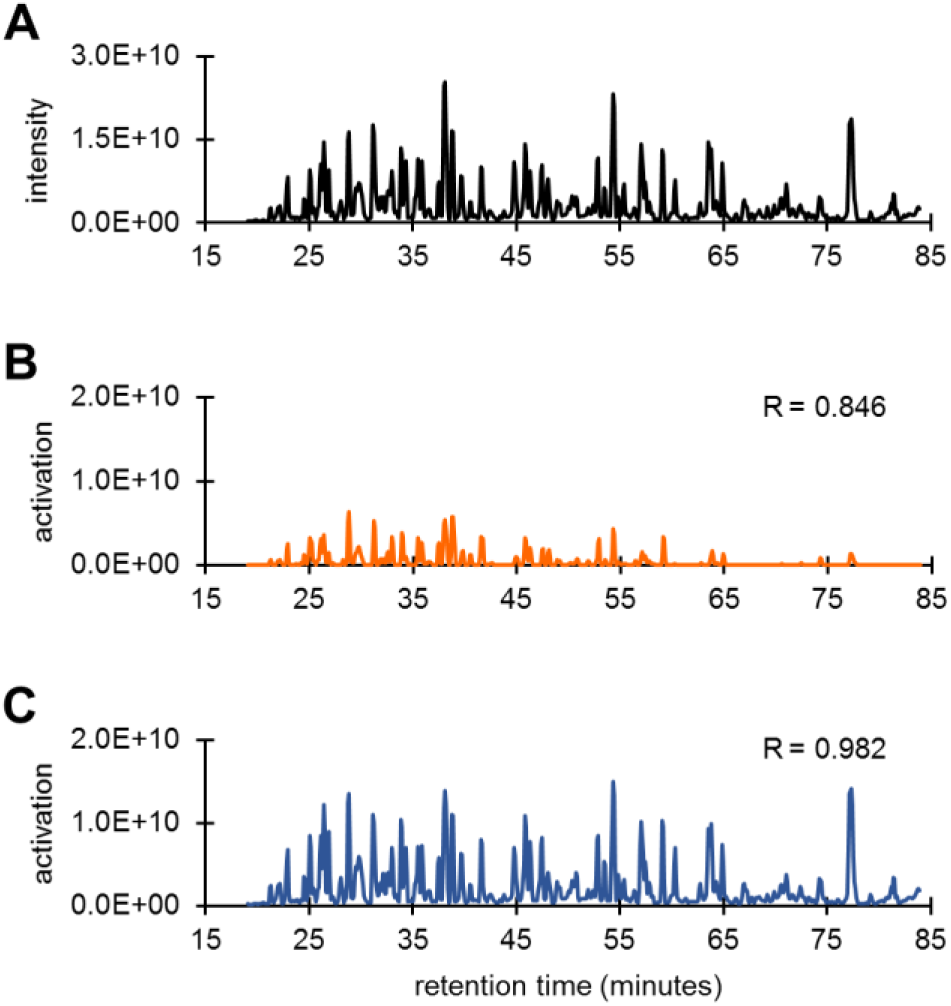
Effects of ProtMSD on chromatogram extraction. (A) The observed total ion current chromatogram. (B) ProtMSD0-approximated chromatogram: the sum of the intensity curves estimated by ProtMSD0. (C) ProtMSD-approximated chromatogram: the sum of the intensity curves estimated by ProtMSD. Correlation coefficients were calculated between the observed total ion currents and the total intensities of each algorithm at the same retention time points.

Next, we compared the retention times of peptides detected by ProtMSD and ProtMSD0 with those estimated by Mascot^10^ and Skyline^11^, two conventional tools used in proteomics. We imported Mascot search results into Skyline MS1-Full Scan Filtering to obtain MS1 parameters. For the peptide ions commonly annotated by ProtMSD and Mascot/Skyline, the retention times detected by ProtMSD0 agreed with those determined by Mascot/Skyline, with *y* = 1.003x, *R* = 0.902 (Figure S1, left), while ProtMSD attained better accuracy, with *y* = 1.000x, *R* = 0.999 (Figure S1, right). These results indicate that our modifications were effective for annotating peptide ions in the mass spectrogram. In order to understand the effects of each type of prior knowledge, we evaluated ProtMSD without using isotopic distribution, retention time initialization, or noise learning (Table 1). The knowledge of isotopic distribution and noise significantly contributed to the recovery of the signal intensity, while retention time information was important for the accuracy of annotation.

**Table 1.** Effects of prior knowledge on the recovery of the signal intensity, the correlation to the observed TIC chromatogram, and the accuracy of peptide retention time annotation

| Prior knowledge |  |  | Performance |  |  |
| --- | --- | --- | --- | --- | --- |
| Noise learning | Isotopic distribution | Retention time | Signal recovery (%) | Chromatogram correlation (R) | Peptide retention time annotation (R) |
| n<sup>a</sup> | n | n | 20% | 0.846 | 0.902 |
| n | y<sup>b</sup> | y | 29% | 0.913 | 0.986 |
| y | n | y | 62% | 0.957 | 0.988 |
| y | y | n | 72% | 0.982 | 0.847 |
| y | y | y | 70% | 0.982 | 0.999 |

a) Without prior knowledge, b): with prior knowledge

#### 1.2 Peptide Identification

ProtMSD uses only MS1 spectra for peptide identification. In contrast, Mascot/Skyline uses MS/MS spectra to acquire sequence-related information, meaning that the success of peptide identification depends to a large extent on the quality of the MS/MS spectra. ProtMSD identified 184 out of 187 (98.40%) peptides identified by Mascot (Figure 3A, left), and the remaining three peptides had 52.29 times lower intensity, 20.78 lower Mascot Peptide Score (PeptScore), and 4.12% lower Skyline isotope dot product (idotp) than the peptides identified in common, calculated by median (Figure S2). In other words, these three peptides had limited intensities in MS1. Moreover, ProtMSD additionally identified 146 peptides, and provided 100% sequence coverage for the four proteins, whereas the conventional method gave only 94.83% coverage on average (Figure S3). Generally, MS1-based methods are more sensitive and can detect more peptides regardless of MS2 information, but MS1 information alone may not be enough for peptide identification if it yields poor profiles, such as distorted isotope patterns.

**Figure 3.**
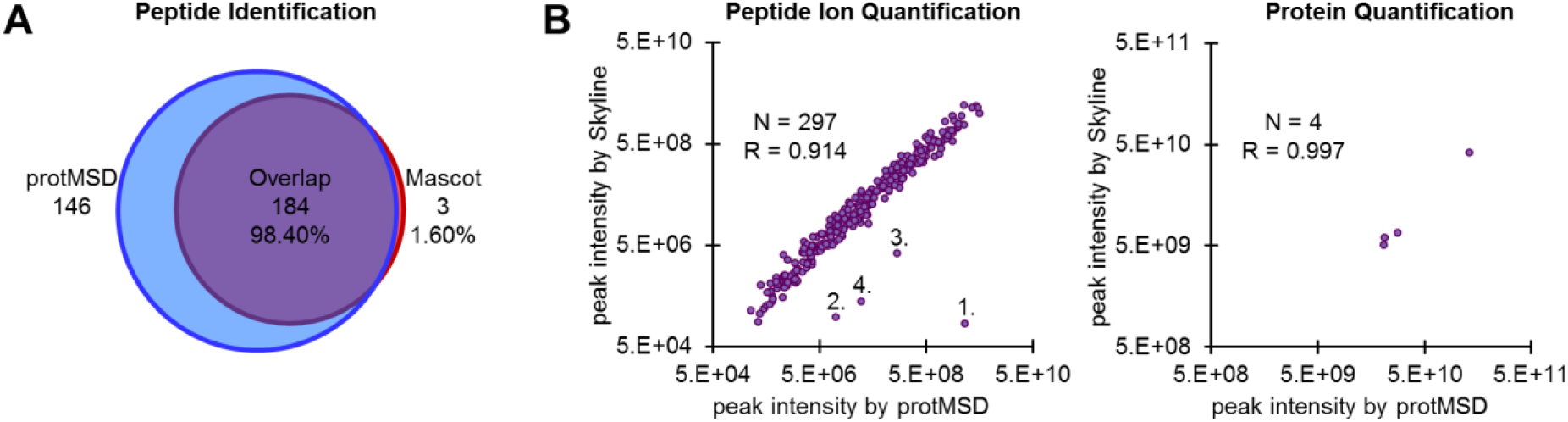
ProtMSD for the four-protein digestion standard. (A) The Venn diagram represents the peptides identified using ProtMSD and Mascot. (B) The peak intensities of peptide ions commonly identified by ProtMSD (x-axis) and Skyline (y-axis) show correlation coefficients of 0.914 with four outliers at the peptide level, and 0.997 at the protein level.

#### 1.3 Protein and Peptide Quantification

We compared the peak intensities obtained by ProtMSD with those obtained by Skyline for commonly identified ions at the peptide ion level. The comparison at the protein level was done using the sum of the intensities of the commonly detected peptide ions. The quantification results for both methods agreed well with each other. Correlation coefficients of 0.997 and 0.914 were obtained at the protein and peptide ion levels, respectively (Figure 3B). We observed four peptide ion outliers calculated from Tukey’s fences, or 1.5 times the interquartile range above the third quartile of the ratio of absolute intensity difference.

We identified two causes of the four outliers, exemplifying the advantage and disadvantage of the algorithms. For outlier 1, Skyline and ProtMSD reported uncorrelated intensities of KLVAASQAALGL with +2 charge for 1.41*e*+05 at 42.1 min and 5.12*e*+09 at 41.60 min. According to Skyline data visualization, this peptide ion had three significant hits at 41.47, 42.10, and 43.13 min by Mascot, but the maximum MS1 intensity was located at 41.61 min, within the dynamic exclusion time. Since Skyline is an MS2-triggered MS1-peak integrated approach and was able to find a small peak close to the significant hit at 42.10 min, it missed the true maximum intensity at 41.61 min because of the short separation distance. Consequently, Skyline underestimated the intensity from the subordinated peak due to dynamic exclusion (Figure S4). In contrast, ProtMSD is a completely MS1-dependent approach, and can report the maximum MS1 peak intensity as an alternative peak quantification tool. Outliers 2, 3, and 4 resulted from HPYFYAPELLFFAKR with charges of +2, +3, and +4. This mixture contained RHPYFYAPELLFFAK, which shares the same precursor isotopic distribution profile as HPYFYAPELLFFAKR; thus, an MS1-based algorithm such as ProtMSD could not distinguish between these two peptides. More features are necessary to quantify peptide isomers uniquely.

To validate the ProtMSD results, we employed a concatenated target-decoy approach^12^ using a shuffle sequence to estimate the false discovery rate (FDR)^13^ as *N*_*D*_/*N*_*T*_, where *N*_*D*_ is the number of decoys and *N*_*T*_ is the number of target-quantified peptides. We also determined the false quantification rate (FQR), calculated as *I*_*D*_/*I*_*T*_, where *I*_*D*_ is the decoy intensity and *I*_*T*_ is the target intensity. ProtMSD gave values of 0.4% FDR and 0.002% FQR.

### 2. Application to *E. coli* proteome analysis

The most frequently used approach to identify peptides is database searching^14^, in which all protein sequences provided in the database are digested *in silico* for reference. An alternative to database search is library search^15^, in which the reference is constructed using a comprehensive collection of confident identification results from previous experiments. Peptide detectability^16^ is a key factor to construct a reference library with suitable size, specificity, and sensitivity for peptide identification and quantification. By using peptide detectability as prior knowledge, library searching can be performed relatively quickly and provides better identification with minimum false positive matches.^17–19^

We examined ProtMSD for analysis of the whole *E. coli* cell lysate with a 70-min gradient. Exploiting the library search concept, we improved ProtMSD by incorporating peptide detectability knowledge. The reference library was the combination of the detected peptides in the *E. coli* cell lysate with a 70-min gradient and with a 550-min gradient, analyzed by Mascot at 1% false discovery rate (FDR) confidence. The dictionary was then constructed to cover all library peptides and extended to include every charge state possible within the *m/z* range. The result from the longer gradient separation provides a larger number of peptides for creating a comprehensive dictionary and offers normalized retention times for initializing an activation matrix. We obtained the normalized retention times by converting the observed retention times to reference retention times based on linear regression with a ±2.5 min window.

#### 2.1 Protein and Peptide Identification

ProtMSD with the peptide library dictionary identified 2877 out of 2998 peptides, or 95.96% of the peptides identified by Mascot (Figure 4A, left). The correlation between the two methods for measuring the retention times of commonly identified peptide ions was high, with y = 0.999x, R = 0.996 (Figure S5). At the protein level, ProtMSD identified more than twice as many proteins as Mascot and included 497 out of 502 proteins (99.00%) identified by Mascot (Figure 4A, right). The conventional method and ProtMSD gave median sequence coverages of 14.80% and 25.12%, respectively (Figure S6).

**Figure 4.**
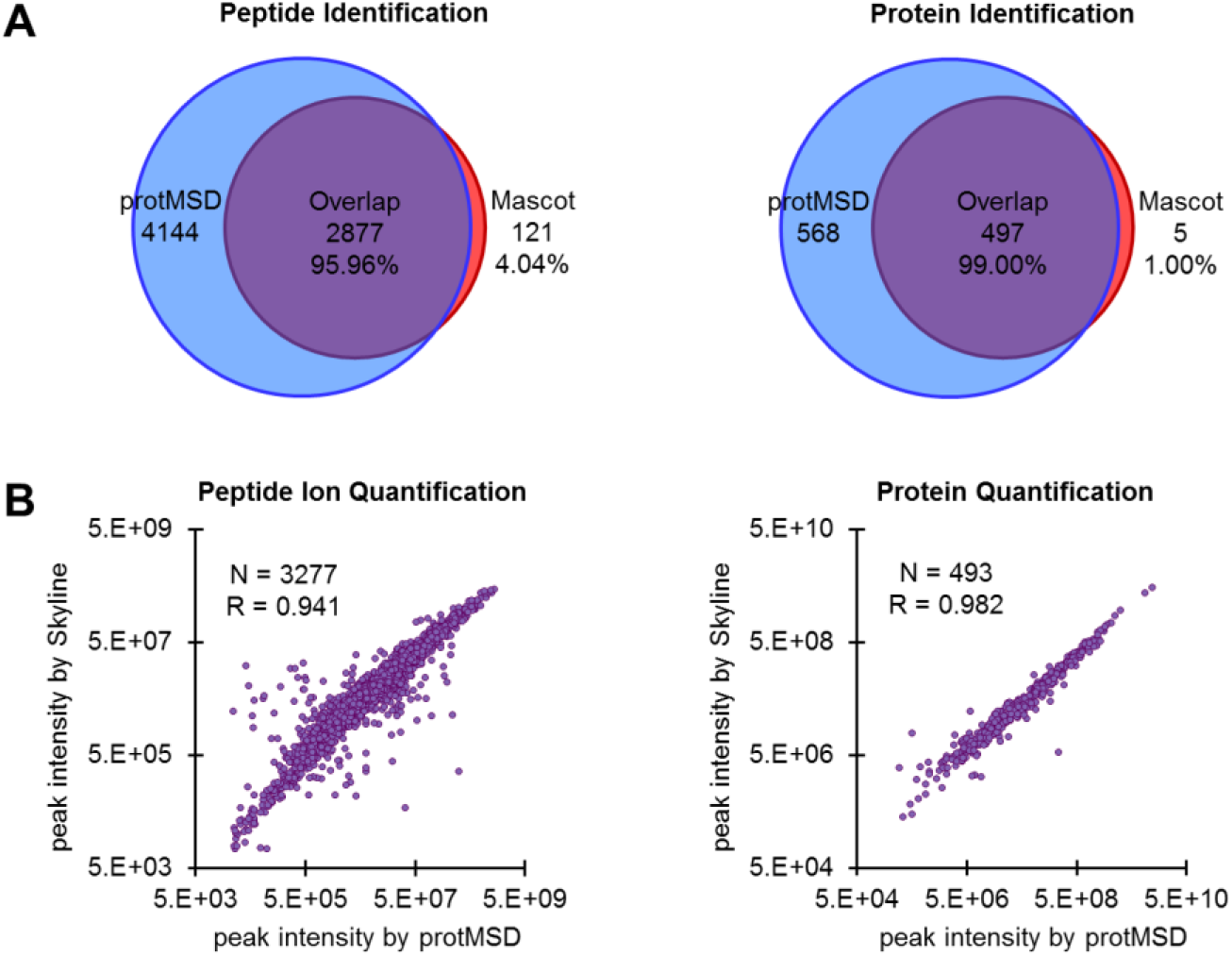
ProtMSD for proteome-scale application. (A) The Venn diagrams represent the E. coli peptides and proteins identified using ProtMSD and Mascot. (B) The peak intensities of peptide ions commonly identified by ProtMSD (x-axis) and Skyline (y-axis) show correlation coefficients of 0.941 at the peptide level and 0.982 at the protein level.

#### 2.2 Protein and Peptide Quantification

Finally, we quantified the peak intensities of commonly identified peptide ions and proteins. After eliminating peaks with S/N ≤ 10, correlation coefficients of 0.982 and 0.941 were obtained at the protein and peptide ion levels, respectively (Figure 4B). We also evaluated the protein quantification based on the peptide ions quantified with each method individually and obtained a correlation coefficient of 0.987 (Figure S7).

Due to the nature of ProtMSD, forcing the well-constructed dictionary to include a large number of false peptides weakens the algorithm performance, leading to the problem of FDR overestimation. In this setting, since the sample used to construct the library and the target sample are the same, FDR in the target sample analysis should be the same as in the library (i.e. 1%). However, in order to validate the power of ProtMSD without the help of any external preprocessing, we also employed a concatenated target-decoy approach. The decoy was generated from frequently identified Hela peptide lists and concatenated to the *E. coli* dictionary with the same number of peptide ions. We obtained 5.4% FDR and 0.5% FQR at the peptide level in this application. The low FQR value indicates that false discovery peaks in ProtMSD are generally small and might be filtered out by appropriate prior knowledge. Further development is needed to accurately estimate the FDR for ProtMSD with a peptide detectability-based library.

### 3. Computational performance

The computational cost of ProtMSD depends on the size of *V* and the transformation of prior knowledge to construct the dictionary or initialize the activation matrix.

For the four-protein digestion mixture, the processing time for ProtMSD0 was 4.50 min. ProtMSD required a longer processing time of 7.30 min due to the extra steps in learning noise, initializing the *H* matrix, and updating the algorithm with a group sparsity constraint. For the *E. coli* proteome sample, ProtMSD with library search took 24.17 minutes for all processing.

## CONCLUSION

We have developed a new statistical method called ProtMSD to identify and quantify peptides and proteins from two-dimensional mass spectrograms obtained in proteomic studies. To formulate ProtMSD, we considered the isotopic distributions of peptides, the mass spectra of noise, and a group sparsity constraint based on the protein-peptide hierarchical relationships, and we incorporated a reasonable initialization based on predicted retention times. The proposed optimization method is guaranteed to converge to a local optimum.

In the analysis of the four-protein digestion mixture, ProtMSD gave a better-resolved chromatogram with a higher accuracy of retention time estimation than its variant without prior knowledge. In the analysis of *E. coli* proteome samples, ProtMSD achieved excellent agreement with the conventional approach based on Mascot/Skyline. In addition, ProtMSD successfully identified more peptides and quantified more than a thousand proteins. The experimental results show that ProtMSD with the library-based dictionary can achieve rapid and large-scale proteome analysis. In this study, we have focused on protein identification and quantification, but the same approach would be readily applicable for the other purposes related to the interpretation of mass spectrograms.

The present results indicate that the matrix decomposition algorithm-based approach is very effective for mass spectrometry-based proteome analysis. Further improvements would be possible by incorporating other significant features, such as product ions from proteomic mass spectrograms. Work along this line is already underway in our laboratory.

## Acknowledgements

This work was supported by the CREST Strategic Basic Research Program from Japan Science and Technology Agency (JST), No. 18070870, and the Database Integration Coordination Program from the National Bioscience Database Center, JST No.18063028.

## Supporting Information Available

The Supporting Information is available free of charge on the ACS Publications website.

Supplementary Figures: S1−S7; Figure S1 Effects of ProtMSD on the accuracy of peptide ion retention time annotation, Figure S2 Profile of peptides exclusively identified by Mascot in the four-protein standard, Figure S3 Sequence coverage comparison between 303 peptide ions identified by Mascot and 596 peptide ions identified by ProtMSD, Figure S4 Outlier investigation for KLVAASQAALGL++ of *m/z* 571.35, Figure S5 Accuracy of peptide ion retention time annotation in proteome-scale application, Figure S6 Sequence coverage comparison between 3509 peptide ions identified by Mascot and 10749 peptide ions identified by ProtMSD, Figure S7 Correlation between *E. coli* proteins quantified by independent ProtMSD and independent Skyline approaches

Supplementary Methods: ProtMSD techniques; Further details of calculation methods, Further details of matrix constructions, ProtMSD algorithm

Experiment setup: Materials, Sample preparation, LC/MS analysis, Mascot/Skyline analysis

(PDF)

Supplementary Table S1-2: Protein and peptide identifications of the four-protein digestion mixture analysis and *E. coli* proteome analysis

(XLSX)

## For Table of Contents Only

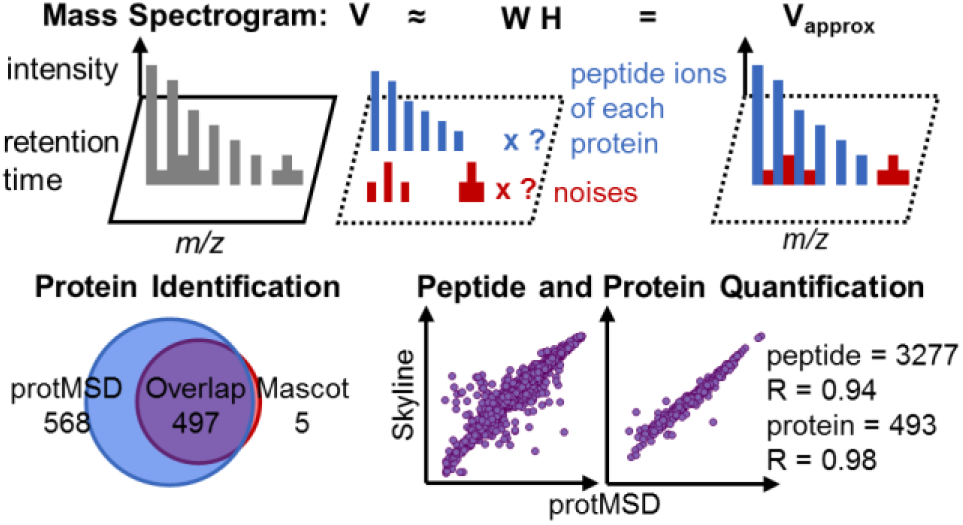

