## Supporting Information for "Peak identification and quantification by proteomic mass spectrogram decomposition" (from another version of this preprint: version 1, 2020-08-06)

**Table of Contents**


- Supplementary Figures
  - Figure S1 Effects of protMSD on the accuracy of peptide retention time annotation.
  - Figure S2 Profile of peptides exclusively identified by Mascot in four-protein standard.
  - Figure S3 Sequence coverage comparison between 303 peptides identified by Mascot and 596 peptides identified by protMSD.
  - Figure S4 Outlier investigation for KLVAASQAALGL++ of *m/z* 571.35.
  - Figure S5 Accuracy of peptide retention time annotation in forty-eight-protein standard.
  - Figure S6 Profile of peptides exclusively identified by Mascot in forty-eight protein standard.
  - Figure S7 Sequence coverage comparison between 1481 peptides identified by Mascot and 3425 peptides identified by protMSD.
  - Figure S8 Accuracy of peptide retention time annotation in proteome-scale application.
  - Figure S9 Sequence coverage comparison between 3509 peptides identified by Mascot and 10749 peptides identified by protMSD
  - Figure S10 The correlation of *E. coli* quantified proteins between independent protMSD and independent Skyline approaches.

- Supplementary Methods
  - protMSD techniques
    - Further details of theory
    - Further details of matrix construction
    - Sparsity constraint weight calculation
    - protMSD algorithm
    - Post-processing
  - Experiment setup
    - Materials

- Sample preparation
- LC/MS analysis
- Mascot/Skyline analysis

### Supplementary Figures

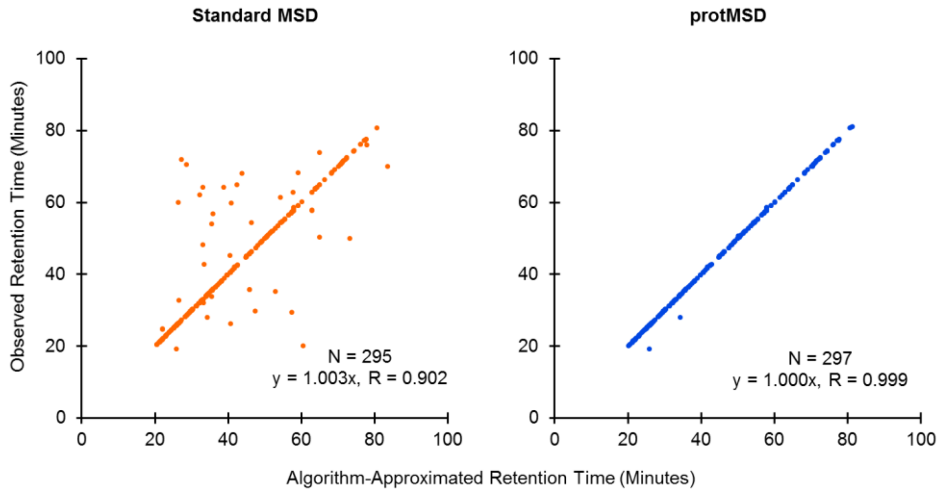


**Figure S1. Effects of protMSD on the accuracy of peptide retention time annotation.**

Proportional relationship between standard MSD (x-axis) and the conventional method (y-axis) for commonly annotated peptide ions (left). Proportional relationship between protMSD (x-axis) and the conventional method (y-axis) for commonly annotated peptide ions (right).

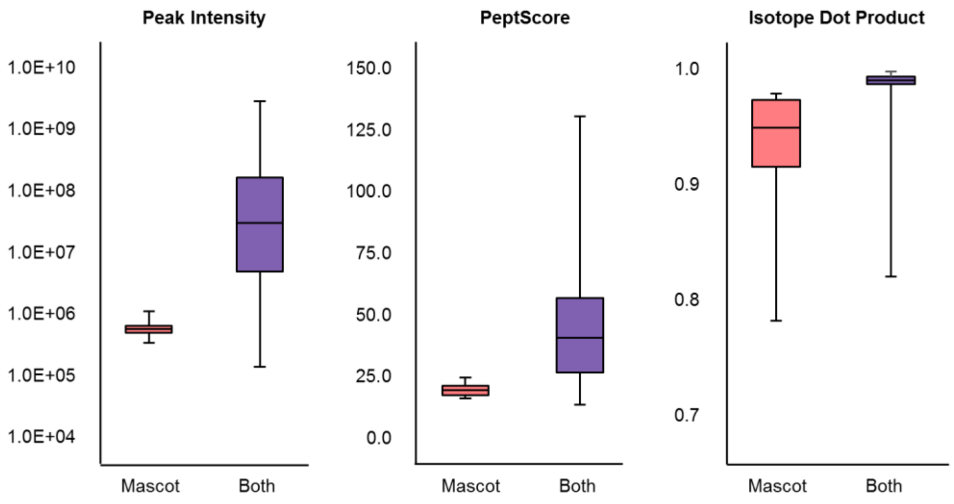


**Figure S2. Profile of peptides exclusively identified by Mascot in four-protein standard.**

The box plot shows that the six peptides exclusively identified by Mascot have 52.29 times lower intensity, 20.78 lower Mascot Peptide Score (PeptScore), and 4.12% lower Skyline isotope dot product (idotp) than the 297 peptides commonly identified by both Mascot and protMSD.

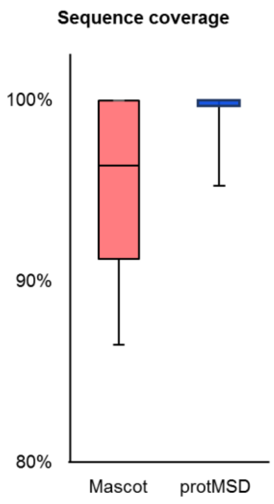


**Figure S3. Sequence coverage comparison between 303 peptides identified by Mascot and 596 peptides identified by protMSD.**

Mascot and protMSD yielded medians of 94.83% and 100.00% coverage, respectively.

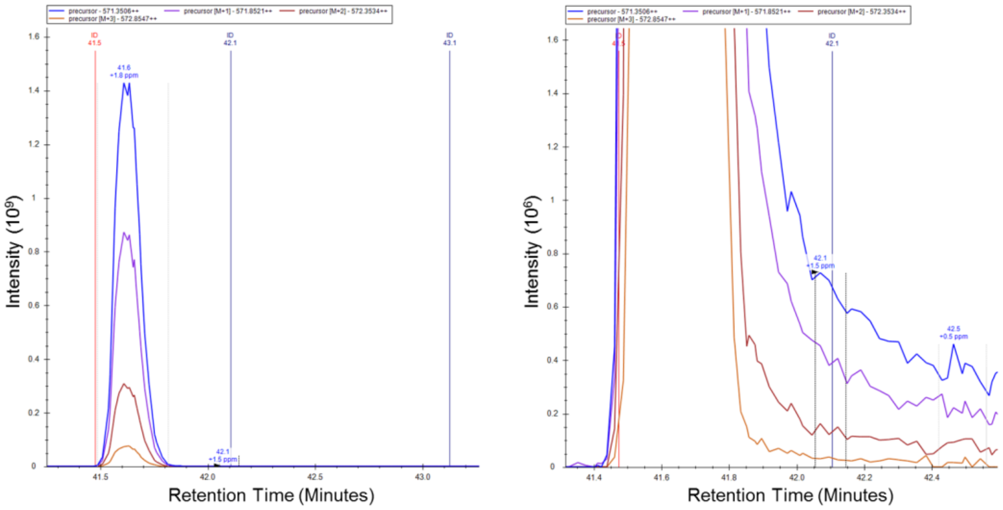


**Figure S4. Outlier investigation for KLVAASQAALGL++ of *m/z* 571.35.**

Skyline graphic interface shows the extracted chromatogram with peak retention time at 41.60 min (left). Detailed examination shows that the Skyline peak integration boundary selecting the peak closest to the significant hit at 42.10 min results in incorrect quantification.

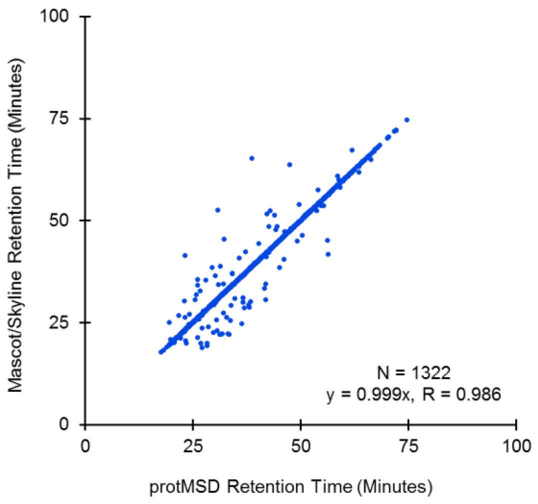


**Figure S5. Accuracy of peptide retention time annotation in forty-eight-protein standard.**

Proportional relationship between protMSD (x-axis) and the conventional method (y-axis) for commonly annotated peptide ions.

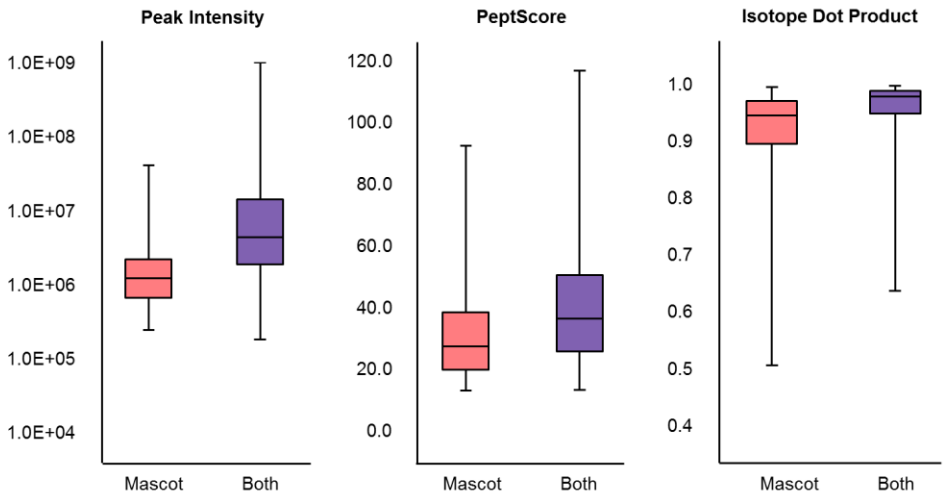


**Figure S6. Profile of peptides exclusively identified by Mascot in forty-eight-protein standard.**

The box plot shows that the 159 peptides exclusively identified by Mascot have 3.45 times lower intensity, 8.59 lower PeptScore, and 3.37% lower idotp than the 1322 peptides commonly identified by both Mascot and protMSD.

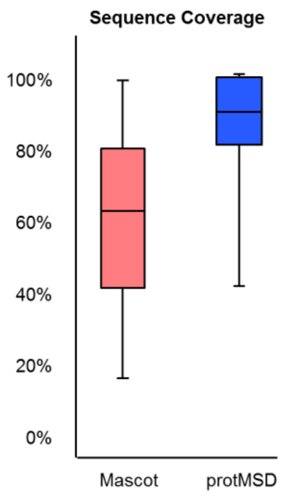


**Figure S7. Sequence coverage comparison between 1481 peptides identified by Mascot and 3425 peptides identified by protMSD.**

Mascot and protMSD yielded medians of 62.02% and 87.29% coverage, respectively.

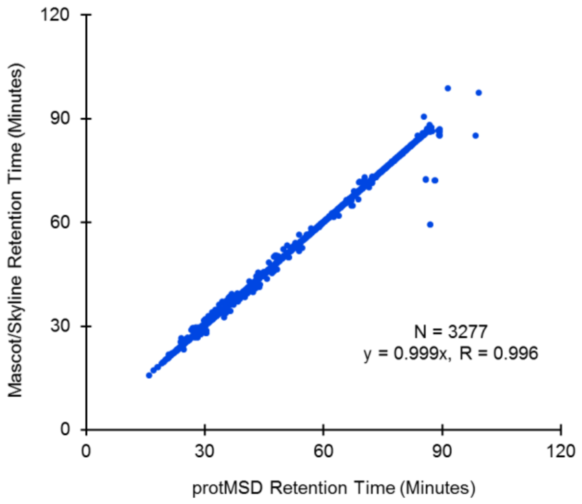


**Figure S8. Accuracy of peptide retention time annotation in proteome-scale application.**

Proportional relationship between protMSD (x-axis) and the conventional method (y-axis) for commonly annotated *E. coli* peptide ions.

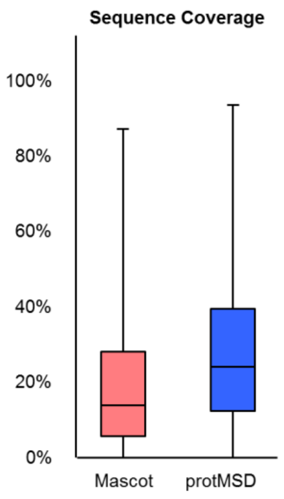


**Figure S9. Sequence coverage comparison between 3509 peptides identified by Mascot and 10749 peptides identified by protMSD.**

Mascot and protMSD yielded medians of 14.80% and 25.12% coverage, respectively.

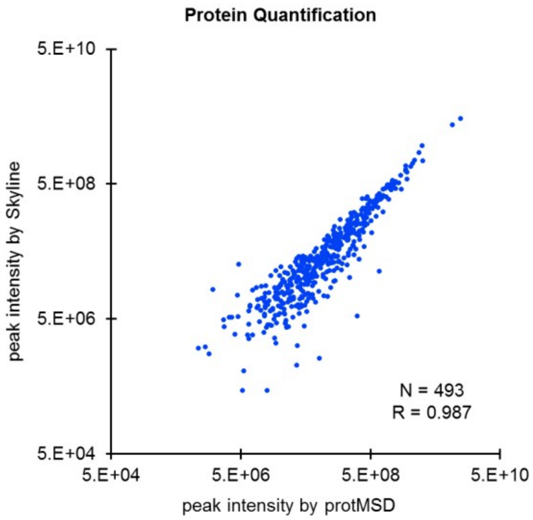


**Figure S10. The correlation of *E. coli* quantified proteins between independent protMSD and independent Skyline approaches.**

The relationship between the sum of all peptide ion peak intensities independently quantified by protMSD (x-axis) and the sum of all peptide ion peak intensities independently quantified by Skyline (y-axis) show correlation coefficients of 0.987.

**Supplementary Methods**

**protMSD techniques**

- **Further details of theory**

  In this work, $D$ is the Kullback-Leibler divergence and $\Omega$ is group sparsity constraint

  - **Kullback-Leibler divergence**

$$D(V \parallel WH) = \sum_{ij}(V_{ij} \log \frac{V_{ij}}{(WH)_{ij}} - V_{ij} + (WH)_{ij})$$

    where $V$ represents the mass spectrogram (*m/z* × retention time), $W$ represents the supervised dictionary of peptide signals (*m/z* × theoretical peptide ion), and $H$ represents the activation results of peptide along the retention time axis (theoretical peptide ion × retention time).

  - **Group sparsity constraint**

$$\Omega = \lambda \log/L1 = \sum_{g=1}^{G} \lambda_g \log(||H_g||_1) + \varepsilon$$

    where $H_g$ is a group of peptide ions digested from the same protein. The majorizing function of $D(V||WH) + \Omega(H_S)$ can be derived for efficient multiplicative updates.[1]

- **Further details of matrix construction**

  - $\boldsymbol{V}$**:** The full scan range of *m/z* and the peptide retention time were divided into fine equal-width bins of 0.01 and 0.005 min, respectively. The retention time was smoothed by a moving average technique using a 12 s time frame and 6 s time frame shift forward.
  - $\boldsymbol{W_S}$**:** The database-based dictionary had *in silico* tryptic digested peptides with up to two missed cleavage sites. Peptide ions ranging in length from seven to 50 amino acids with charges ranging from +2 to +4 were included. Cysteine carbamidomethylation was set as a fixed modification. Methionine oxidation and N-terminal cysteine ammonia loss were set as variable modifications. Features of six isotopic peaks including the monoisotopic peak were calculated, normalized, and assigned to columns of $W_S$. The library-based dictionary had Mascot-identified peptide ions and the additional peptide ions for all possible charges in *m/z* range. Cysteine carbamidomethylation and methionine oxidation were set as fixed and variable modifications, respectively. Features of six isotopic

peaks including the monoisotopic peak were calculated, normalized, and assigned to columns of $W_S$.

- $\boldsymbol{W_N}$**:** Noise was learned from a blank region of the mass spectrogram, i.e., the last 10 min of gradient elution, which was free from peptides, by using scikit-learn[2] NMF with *k* = 2. The obtained patterns were assigned to two columns of $W_N$.
- $\boldsymbol{H_S}$**:** The activations of each peptide were restricted to zero at retention time frames in which the peptide peaks were highly unlikely to be detected.
  - For protMSD with the database-based dictionary, two initializations were combined using the predicted retention times and the similarities to the theoretical isotopic distribution, respectively. The predicted retention times were obtained from the algorithm, achrom[3], which could be calibrated by using an in-house data set. The points within the possible retention time frame were initialized using Gaussian distribution ($\mu$ = the predicted retention time, $\sigma$ = the average peak width/4). The similarities to the theoretical isotopic distribution were calculated for peptide ions by using cosine similarity (*cos*) at *m/z* $i$ and time $j$ as

    $$cos = \frac{\mathcal{W}_x \cdot \mathcal{V}_x}{\|\mathcal{W}_x\| \; \|\mathcal{V}_x\|}$$

    where $\mathcal{W}_x$ is peaks in $W_i$ and $\mathcal{V}_x$ is peaks at $x$ in $V_j$

    In order to include the surrounding distributions, *cos* were calculated at $j-1$, $j+1$, and $x'$ which included the previous isotopic index into $x$. All *cos* were summed and calculated for average of median and mean. In this work, the points outside 25-min frame and the *cos* less than 0.5 were restricted to zero. Wide possible retention time frame and low *cos* were set to exclude only highly unreasonable retention times, and they could be adjusted according to the experiment.
  - For protMSD with the library-based dictionary, the predicted retention times were collected from references by combining between Mascot peak retention times from a 70-min gradient experiment and converted retention times of a 550-min gradient experiment using linear regression. Although Mascot-identified retention times are triggered by MS2, they are very dependable. The possible retention time were set for 5 minutes and *cos* was not used.

- $\boldsymbol{H_N}$**:** The points in the peptide subspace were fully initialized with positive values.

- **Sparsity constraint weight calculation**

  The sparseness measure[4] with protein size $\boldsymbol{\lambda} = [\boldsymbol{\lambda_1}, \boldsymbol{\lambda_2}, \dots, \boldsymbol{\lambda_G}]^T$ can be calculated as follows

$$\boldsymbol{\lambda} = \frac{1}{\sqrt{m}} \sum_{i}^{m} \frac{\sqrt{t} - ||V_i||_1/||V_i||_2}{\sqrt{m-1}} \boldsymbol{p}$$

  where the $V$ matrix has $m$ bins of *m*/*z* and $t$ frames of retention time ($m \times t$), and the $\boldsymbol{p}$ vector represents the number of peptide ions for each protein group as $\boldsymbol{p} = [\boldsymbol{p_1}, \boldsymbol{p_2}, \dots, \boldsymbol{p_G}]^T$. In the case of protMSD with the library-based dictionary, since the activations are more restricted than protMSD with the database-based dictionary, a window-gradient ratio is incorporated by multiplying window (minute)/gradient (minute) to $\boldsymbol{\lambda}$.

- **protMSD algorithm**

  Since protMSD has a supervised dictionary, the algorithm aims to update $H$ by iterating between:

  1. updating $H$
  2. updating $H_g$
  3. checking $D(V||WH) + \Omega(H_S)$. If the change since the last iteration is less than the tolerance, the algorithm declares convergence.

The algorithm is summarized below:

**Algorithm 1**: Modified NMF

**input** : $V, W = [W_S \;\; W_N]$
**initialize** : $H$
**repeat**
    # updating $H$
    $H \leftarrow H .* (W^T(V./WH))$;
    # updating $H_g$
    **for** $g = 1{:}G$ **do**
        $H_g \leftarrow \frac{1}{1 + \lambda_g/(||H_g||_1 + \varepsilon)} H_g$;
    **end**
**until** *convergence*;
**return** $H$

where $.*$ and ./ are component-wise multiplication and division, and $(.)^T$ is matrix transposition.

- **Post-processing**

  For peptide identification, the criterion was peak activated intensity with S/N > 3. For peptide quantification, the criterion was peak activated intensity with S/N > 10. For protein identification, at least two unique identified peptides were required per protein.

  The highest activation peak with cosine similarity more than 0.90 was selected as a default candidate. In the design of using the database-based dictionary, the additional post-processing was performed by evaluating the other two highest annotated peaks for each peptide ion. If one of them has cosine similarity more than 0.97 and 20% better than the others, the default candidate will be replaced by the better peak.

The mass spectrograms from different instruments and proteomic experiments have different features. In order to obtain an optimal result from protMSD, we strongly suggest that the parameters should be carefully set for each observed mass spectrogram.

### Experiment setup

- **Materials:**

  Recombinant human albumin, catalase from erythrocytes, recombinant human epidermal growth factor, recombinant human leptin, and universal proteomic standard (UPS1) were obtained from Sigma-Aldrich (Darmstadt, Germany). Dithiothreitol, iodoacetamide, Lys-C, trifluoroacetic acid (TFA), acetronitrile (ACN), and piperidine were obtained from Wako (Osaka, Japan). Trypsin was obtained from Promega (Madison, WI).

- **Sample preparation:**

  For four-protein standard: 25 ng of albumin, catalase, EGF, and leptin were mixed. The sample was reduced with dithiothreitol (10% v/v) and alkylated with iodoacetamide (10% v/v) before Lys-C and trypsin digestion (1% w/w) for 3 h and overnight, respectively. Peptides were desalted with StageTips[5] and suspended in the loading buffer (0.5% TFA and 4% ACN) for subsequent LC/MS analysis.

  For forty-eight-protein standard: a commercial UPS1 was premixed with 5 pmol of 48 proteins ranging in molecular mass from 6000 to 83,000 daltons. The processes of reduction, alkylation, digestion, and desalting for subsequent LC/MS analysis were the same as described for the previous sample.

- **LC/MS analysis**

  A self-pulled analytical column was prepared with ReproSil-Pur C18-AQ materials (Dr. Maisch, Ammerbuch, Germany). The mobile phases consisted of (A) 0.5% acetic acid and (B) 0.5% acetic acid in 80% ACN. A gradient condition with flow rate 500 nL/min was set: 5–10% B in 5 min, 10–40% B in 60 min, 40–100% B in 5 min, 100% B for 10 min, and 5% B for 30 min. An Ultimate 3000 pump (Thermo Fisher Scientific, Germering, Germany) and a HTC-PAL autosampler (CTC Analytics, Zwingen, Switzerland) were used coupled to a Q-Exactive hybrid quadrupole-orbitrap mass spectrometer (Thermo-Fisher Scientific). Tandem mass spectra were acquired using data-dependent acquisition. The MS1 survey scan range was scanned at a resolution of 70,000 for *m/z* 300–1500, an AGC level of 3*e*+6, and a maximum ion accumulation time of 100 ms. The top 10 ions having charges from +2 to +8 were selected in a scan range of *m/z* 200–2000 with an isolation window of *m/z* 2. The

AGC target was $1e+5$, and the maximum ion accumulation time was 100 ms. A dynamic exclusion with a duration of 30 s was used to reduce repeated ions.

- **Mascot[6]/Skyline[7] analysis**

**Software version**

| | | |
|---|---|---|
| Mascot version | : | 2.7.0 |
| Database | : | Ecoli_K12_all 20180721 |
| Skyline version | : | 4.2.0. 19072 (64-bit) |

**Search parameters**

| | | |
|---|---|---|
| Enzyme | : | Lys-C/Trypsin |
| Fixed modifications | : | Carbamidomethyl (C) |
| Variable modifications | : | Ammonia-loss (N C-term), Oxidation (M) |
| Peptide mass tolerance | : | ± 10 ppm |
| Fragment mass tolerance | : | ± 20 ppm |
| Peptide charges | : | +2, +3, +4 |
| Max missed cleavages | : | 2 |
